# A noncanonical FLT3 gatekeeper mutation disrupts gilteritinib binding and confers resistance

**DOI:** 10.1101/2021.01.27.428336

**Authors:** Sunil K. Joshi, Setareh Sharzehi, Janét Pittsenbarger, Daniel Bottomly, Cristina E. Tognon, Shannon K. McWeeney, Brian J. Druker, Elie Traer

**Author notes:** **CORRESPONDENCE:** Elie Traer, M.D., Ph.D.; Oregon Health & Science University, 3181 SW Sam Jackson Park Road, Mail Code: KR-HEM, Portland, OR 97239.

## Abstract

The recent FDA approval of the FLT3 inhibitor, gilteritinib, for AML represents a major breakthrough for treatment of FLT3 mutated AML. However, patients only respond to gilteritinib for 6-7 months due to the emergence of drug resistance. Clinical resistance to gilteritinib is often associated with expansion of NRAS mutations, and less commonly via gatekeeper mutations in FLT3, with F691L being the most common. We developed an *in vitro* model that charts the temporal evolution of resistance to gilteritinib from early microenvironmental-mediated resistance to late intrinsic resistance mutations. Our model system accurately recapitulates the expansion of NRAS mutations and the F691L gatekeeper mutations found in AML patients. As part of this study, we also identified a novel FLT3^N701K^ mutation that also appeared to promote resistance to gilteritinib. Using the Ba/F3 system, we demonstrate that N701K mutations effectively act like a gatekeeper mutation and block gilteritinib from binding to FLT3, thereby promoting resistance. Structural modeling of FLT3 reveals how N701K, and other reported gilteritinib resistance mutations, obstruct the gilteritinib binding pocket on FLT3. Interestingly, FLT3^N701K^ does not block quizartinib binding, suggesting that FLT3^N701K^ mutations are more specific for type 1 FLT3 inhibitors (gilteritinib, midostaurin, and crenolanib). Thus, our data suggests that for the FLT3^N701K^ mutation, switching classes of FLT3 inhibitors may restore clinical response. As the use of gilteritinib expands in the clinic, this information will become critical to define clinical strategies to manage gilteritinib resistance.

## INTRODUCTION

Acute myeloid leukemia (AML) is a genetically heterogenous disease with approximately 20,000 new cases per year in the United States^1, 2^. Patients with AML have a 5-year survival of <25%, and intense efforts are underway to develop new treatments to improve survival^1^. Mutations in the FMS-like tyrosine kinase-3 (FLT3) gene are among the most common genomic aberrations in AML. Internal tandem duplication (ITD) in the juxtamembrane domain of FLT3 are present in approximately 20% of patients with AML. These mutations cause constitutive kinase activity, and lead to an increased risk of relapse and reduced survival. Another set of mutations in the tyrosine kinase domain (TKD) of FLT3 occur in 5-10% of AML patients. In contrast to FLT3-ITD, FLT3 TKD mutations result in less activation of FLT3 and do not increase the risk of relapse^3^.

Multiple FLT3 tyrosine kinase inhibitors have been developed and can be separated into two classes. Type I inhibitors are canonical ATP competitors that bind the ATP binding site of FLT3 in the active conformation and are effective against both ITD and TKD mutations. By contrast, type II inhibitors bind the hydrophobic region adjacent to the ATP binding domain in the inactive conformation. Type II inhibitors are effective against FLT3-ITD, but do not inhibit FLT3 TKD mutations. Quizartinib, a type II inhibitor, has potent activity against FLT3, KIT, and RET. Despite high response rates as a monotherapy in patients with relapsed/refractory AML, the duration of response to quizartinib is approximately 4 months, and resistance via FLT3 TKD mutations is common^4–6^. These mutations occur frequently at the activation loop residue D835 and less commonly at F691 which represents the “gatekeeper” position in FLT3^4^.

Gilteritinib is second-generation inhibitor that targets FLT3 and AXL^7^. As a type I inhibitor, it is active against TKD mutations that impart quizartinib resistance. It was approved as monotherapy in relapsed/refractory patients with AML based upon the randomized phase 3 clinical study (ADMIRAL) which compared gilteritinib with chemotherapy^7^. Despite the significant survival benefit in the gilteritinib arm, monotherapy is limited by the development of resistance, which typically occurs after 6-7 months. Resistance to gilteritinib most commonly occurs through acquisition/expansion of NRAS mutations, however a minority of patients with F691L gatekeeper mutations were also identified^8^. To search for additional resistance mutations to gilteritinib, Tarver *et al.* used a well-established ENU mutagenesis assay and identified Y693C/N and G697S as mutations that confer resistance *in vitro*^6^. These mutations appear to function similar to the gatekeeper mutation by blocking gilteritinib binding to FLT3, but have not been reported in patients.

## RESULTS & DISCUSSION

To more broadly investigate mechanisms of resistance to gilteritinib, we developed a two-step model of resistance that recapitulates the role of the marrow microenvironment (**Figure 1A**). In the first stage of resistance, or early resistance, the FLT3-mutated AML cell lines MOLM14 and MV4;11 are cultured with exogenous ligands, fibroblast growth factor 2 (FGF2) and FLT3 ligand (FL), that are normally supplied by marrow stromal cells. These culture conditions allow the cells to become resistant to gilteritinib without the need for resistance mutations^9^. When ligands are removed, the cells regain sensitivity to gilteritinib, but ultimately become resistant, which we term late resistance. At this point, intrinsic resistance mutations were identified in all of the cultures via whole exome sequencing. Similar to clinical data^7^, we found that the most common mutations are activating mutations in NRAS^10^. One late resistant culture had an FLT3^F691L^ gatekeeper mutation, and 3 cultures had an FLT3^N701K^ mutation, which has not previously been reported (**Figure 1B**). Given its proximity to F691L (**Figure 1C-D**), we hypothesized that this mutation might also disrupt gilteritinib binding to FLT3.

**Figure 1:**
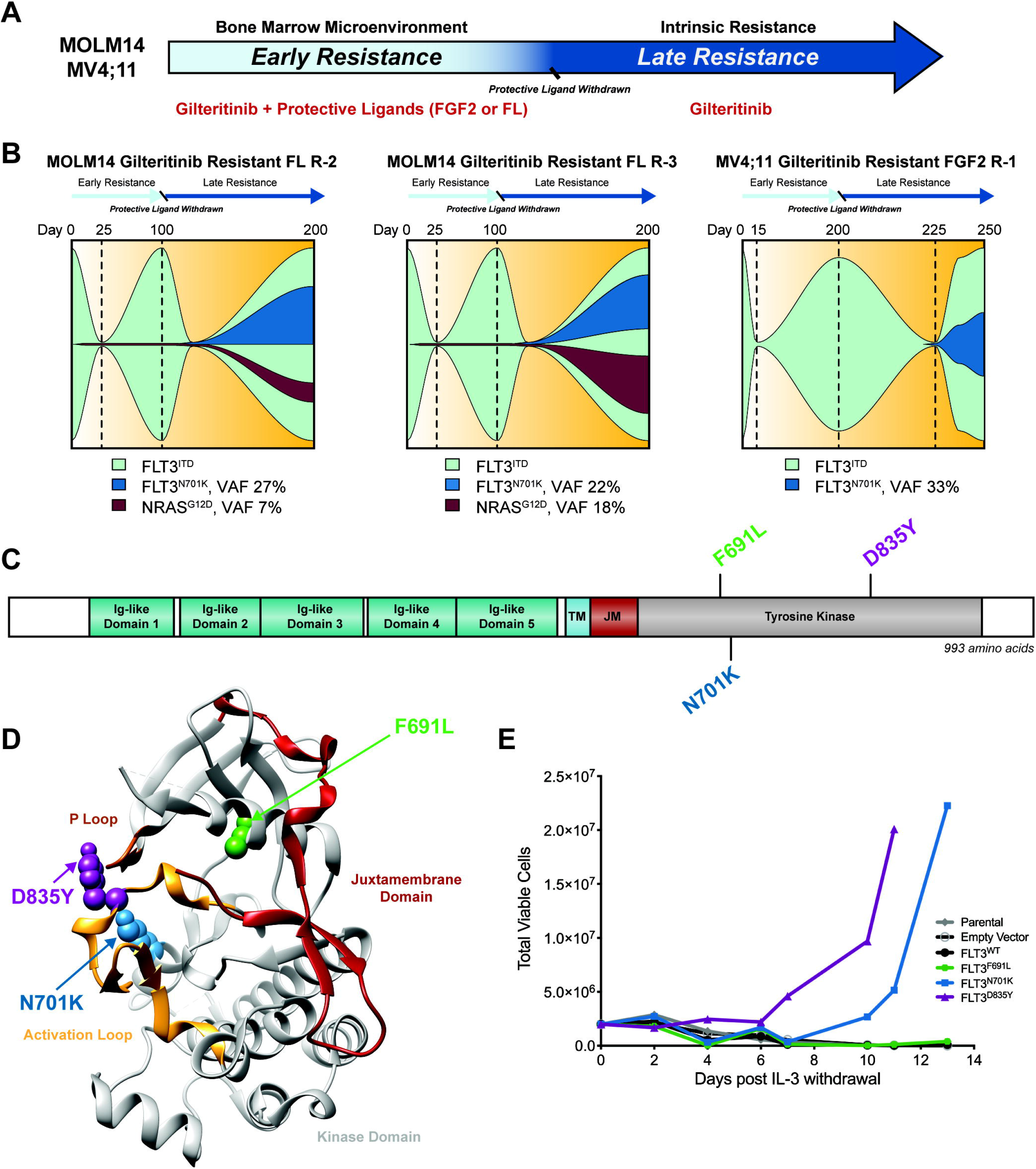
FLT3^N701K^ is an oncogenic mutation. **A.** Model of early and late gilteritinib resistance. MOLM14 (N = 8) and MV4;11 (N = 8) cultures were continuously treated with gilteritinib and exogenous microenvironmental ligands (FGF2 or FL, 10 ng/mL) to recapitulate the role of the marrow microenvironment in the development of early resistance. Following ligand withdrawal, cultures become transiently sensitive to gilteritinib again, but eventually become resistant with the outgrowth of NRAS and FLT3 resistance mutations that drive late resistance. **B.** Potential clonal evolution paths resulting in the outgrowth of the FLT3^N701K^ mutation in three cultures. Mutations were identified by whole exome sequencing and displayed via fishplots^14^. All mutations were confirmed by Sanger sequencing. **C.** Gene schematic depicts location of the FLT3^N701K^ point mutation relative to FLT3 gatekeeper (F691L) and activating loop (D835Y) mutations. The location of the following domains is included: immunoglobulin (Ig)-like loops, transmembrane (TM), juxtamembrane (JM), and tyrosine kinase. **D.** Ribbon diagram mapping the location of FLT3^N701K^ onto the crystal structure of the FLT3 kinase domain. Diagram was adapted from PDB 1RJB^3^ and visualized with the UCSF Chimera software^15^. **E.** FLT3^N701K^ transforms the murine Ba/F3 pro-B cell line and enables IL-3 independent growth. No growth was observed in parental Ba/F3 cells or cells harboring an empty vector (pMX-puro), wildtype (WT) FLT3, or FLT3^F691L^. Total viable cells are plotted over time and cell growth was measured after the withdrawal of IL-3. This experiment was repeated at least twice with consistent results.

To determine whether the FLT3^N701K^ mutation has oncogenic capacity, we evaluated this mutation in the Ba/F3 transformation assay. Ba/F3 cells are normally IL-3 dependent but the presence of certain oncogenes transforms them to grow indefinitely in the absence of IL-3^11^. The FLT3^N701K^ mutation, similar to FLT3^ITD^ and FLT3^D835Y^, is an activating mutation and promoted growth of Ba/F3 cells in the absence of IL-3, whereas the parental, empty vector, FLT3 wild type (FLT3^WT^), or FLT3^F691L^ did not confer IL-3-independent growth (**Figure 1E**).

In contrast to Ba/F3 cells expressing FLT3^D835Y^, Ba/F3 cells with FLT3^N701K^ were much less sensitive to gilteritinib with an approximate 8.5-fold increase in IC_50_ (**Figure 2A**). To test whether FLT3^N701K^ also promoted resistance to gilteritinib in the presence of FLT3^ITD^ mutations (**Figure 1B**), we generated FLT3^ITD + N701K^ and FLT3^ITD + F691L^ double mutants and expressed them in Ba/F3 cells. Concordant with previous studies^4^, the FLT3^ITD + F691L^ mutant demonstrated an approximate 11-fold increase in IC_50_ to gilteritinib compared to FLT3-ITD alone. FLT3^ITD+N701K^ Ba/F3 cells were nearly identical to FLT3^ITD + F691L^ cells in their resistance to gilteritinib (**Figure 2B**). As a control, FLT3^WT^ Ba/F3 cells grown with IL-3 were insensitive to gilteritinib at comparable doses.

**Figure 2:**
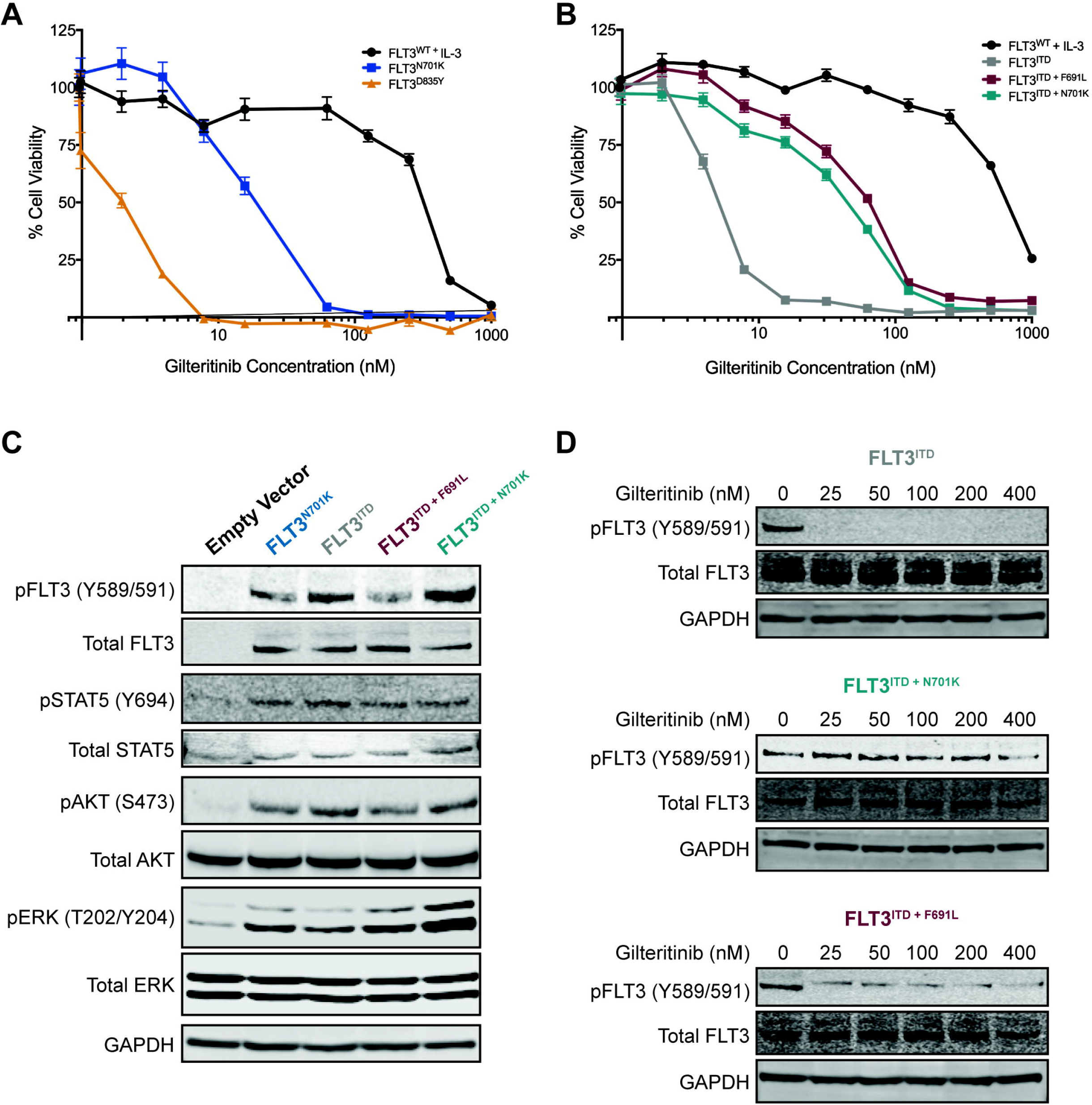
FLT3^N701K^ confers resistance to gilteritinib. **A-B.** Ba/F3 cells expressing FLT3^N701K^ and FLT3^ITD+N701K^ demonstrate reduced gilteritinib sensitivity. FLT3^D835Y^ and FLT3^ITD+F691L^ were used as historical controls. Six replicates of WT and mutant FLT3 Ba/F3 cells were plated with a dose gradient of gilteritinib (0 – 1000 nM) for 72 hrs. FLT3^WT^ cells were plated in media supplemented with IL-3. Cell viability was determined using a tetrazolamine-based viability assay. Viability is represented as a percentage of the untreated control. The average mean ± SEM is shown. **C.** Expression of total and phosphorylated FLT3 is increased in mutant-transformed Ba/F3 cells relative to cells harboring empty vector. All mutants phosphorylate canonical downstream effectors – STAT5, AKT, and ERK. GAPDH served as a loading control. Prior to lysis, empty vector cells were grown in IL-3 supplemented media and all lines were starved overnight in 0.1% BSA RPMI. **D.** FLT3 activity is sustained with mutations in N701K and F691L. Ba/F3 cells harboring FLT3^ITD^, FLT3^ITD+F701K^, and FLT3^ITD+F691L^ were treated with gilteritinib (0 – 400 nM) for 90 minutes and lysed for immunoblot analysis^4, 6^.

Next, we assessed the impact of FLT3^N701K^ mutations on downstream FLT3 signaling pathways. Ba/F3 cells transformed with FLT3^N701K^, FLT3^ITD^, FLT3^ITD + F691L^, and FLT3^ITD + N701K^ all resulted in phosphorylation of FLT3 (Y589/591) and STAT5 (Y694), AKT (S473), and ERK (T202/Y204) (**Figure 2C**). However, only FLT3^ITD + N701K^ or FLT3^ITD + F691L^ showed sustained phospho-FLT3 with increasing concentrations of gilteritinib (**Figure 2D**), indicating that both of these mutations prevent gilteritinib inhibition of FLT3, particularly at lower doses. The FLT3 kinase activity as reflected by FLT3 phosphorylation mirrored the viability assays in **Figure 2B**.

Since F691L gatekeeper mutations are known to drive resistance to multiple FLT3 inhibitors^4, 8, 12, 13^, we treated FLT3^ITD^, FLT3^ITD + N701K^ and FLT3^ITD + F691L^ Ba/F3 cells with midostaurin, crenolanib, and quizartinib. Although FLT3^ITD + F691L^ and FLT3^ITD + N701K^ were largely insensitive to type I inhibitors midostaurin and crenolanib, cells with FLT3^ITD + N701K^ were notably more sensitive to the type II inhibitor quizartinib (**Supplemental Figure 1**), suggesting that N701K blocks gilteritinib binding of type I inhibitors more effectively than type II. This was further apparent from our modeling of the FLT3^N701K^ mutation. While the FLT3^N701K^ mutation may sterically interfere with the binding of gilteritinib, quizartinib binding does not appear to be affected (**Supplemental Figure 2**).

Through our studies, we identified the novel FLT3^N701K^ mutation in addition to the FLT3^F691L^ gatekeeper mutation. We used the Ba/F3 system to demonstrate that N701K blocks gilteritinib binding to FLT3, similar to the gatekeeper F691L, and promotes resistance to gilteritinib. Our data fit nicely with recent data from a mutagenesis screen of Ba/F3 cells with FLT3-ITD that identified F691L in addition to D698N, G697S, and Y693C/N as mutations that drive resistance to gilteritinib^6^. Modeling of these mutations indicates that they cause the loss of hydrogen bonding that accommodates the FLT3 side chain, leading to a steric clash between the tetrahydropyran ring of gilteritinib and FLT3^6^. Given the proximity of N701K to these mutations, we speculate that the mechanism of resistance to gilteritinib imparted by this mutation is similar (**Supplemental Figure 2**). Importantly, these complementary methods identify a common hotspot for gilteritinib resistance mutations (**Supplemental Figure 3**). Given the increasing use of gilteritinib in the clinic, we anticipate that additional resistance mutations will likely be identified in patients. Of note, the N701K mutation appears to be more resistant to type I inhibitors but retains sensitivity to type II inhibitors such as quizartinib (**Supplemental Figure 1**), implicating that TKI class switching could serve as a promising avenue to mitigate development of gilteritinib resistance. The use of type I FLT3 inhibitors following the acquisition of resistance to type II inhibitors is a well-established approach to overcome resistance. However, what makes the case with the N701K mutation interesting is acquired sensitivity to a type II inhibitor following development of resistance to a type I inhibitor, which is a largely underappreciated concept. This knowledge can be used to help rationally sequence FLT3 inhibitors upon development of resistance.

## ACKNOWLEDGEMENTS

We are extremely thankful to the Massively Parallel Sequencing Shared Resource for technical support. We thank Sudarshan Iyer for assistance with UCSF Chimera. This work was supported by the American Cancer Society (MRSG-17-040-01-LIB) to E.T., the Drug Sensitivity and Resistance Network (DRSN; U54CA224019) to B.J.D, S.K.M, and E.T, and Cancer Center Support grant (5P30CA069533-22) to B.J.D. Additional funding was provided by the Howard Hughes Medical Institute, and The Leukemia & Lymphoma Society to E.T. and B.J.D. S.K.J. is supported by the ARCS Scholar Foundation, The Paul & Daisy Soros Fellowship, and the National Cancer Institute (F30CA239335). E.T. is supported by grants from the Leukemia & Lymphoma Society, Hildegard Lamfrom Physician Scientist Award, American Cancer Society, and Cancer Early Detection Advanced Research Center. C.T., D.B., S.K.M., B.J.D, and E.T. were all supported by National Cancer Institute (U54CA224019).

## AUTHOR CONTRIBUTIONS

**Study supervision:** E. Traer

**Conception and design:** S.K. Joshi, E. Traer

**Development of methodology:** S.K. Joshi, E. Traer

**Acquisition of data:** S.K. Joshi, S. Sharzehi, J. Pittsenbarger

**Analysis and interpretation of data (e.g., statistical analysis, biostatistics, computational analysis):** S.K. Joshi, S. Sharzehi, J. Pittsenbarger, D. Bottomly, C.E. Tognon, S.K. McWeeney, B.J. Druker, E. Traer

**Writing, review, & editing of the manuscript:** S.K. Joshi, J. Pittsenbarger, S. Sharzehi, D. Bottomly, C.E. Tognon, S.K. McWeeney, B.J. Druker, E. Traer

**Development of prioritization framework to assist in rigor and reproducibility:** D. Bottomly, S.K. McWeeney

---

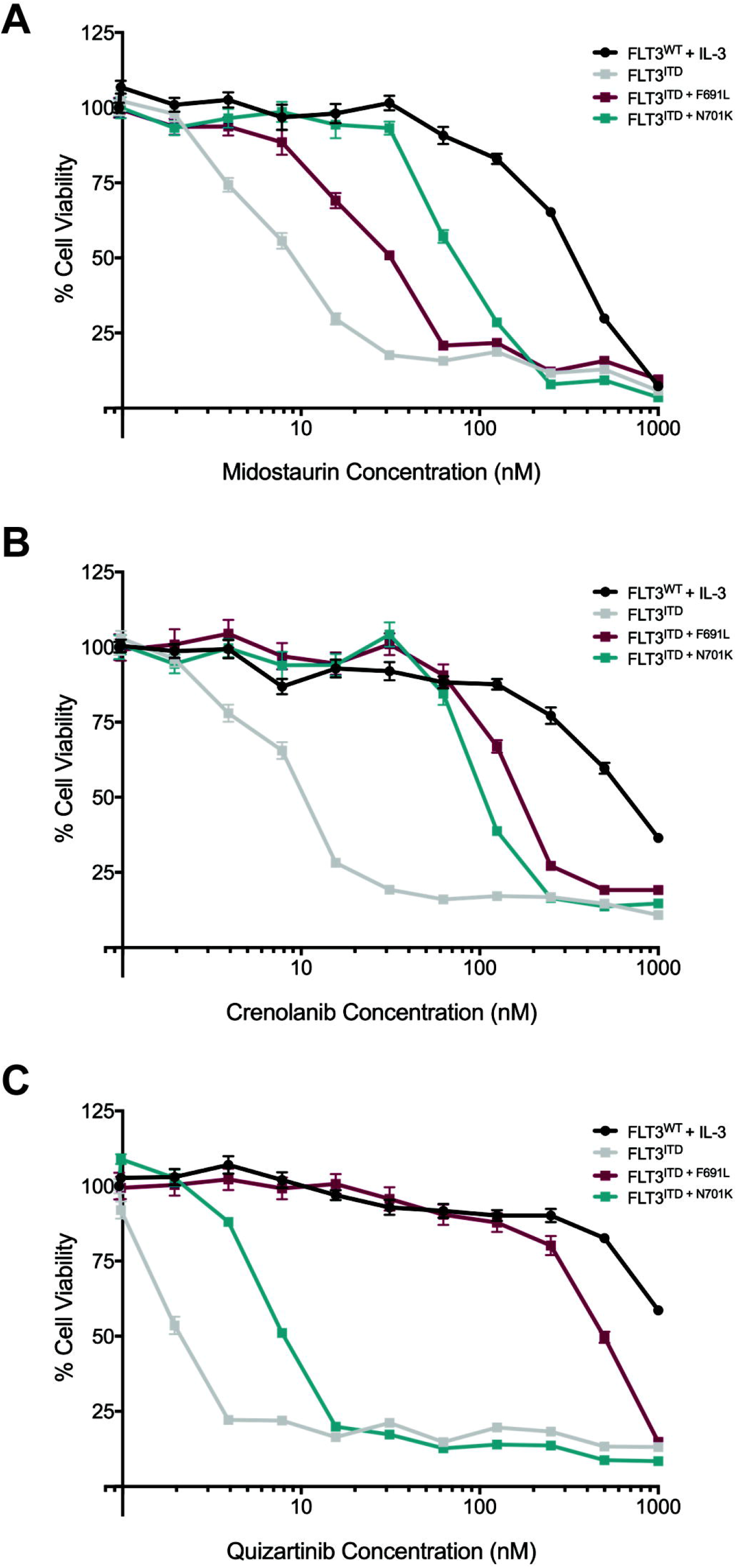

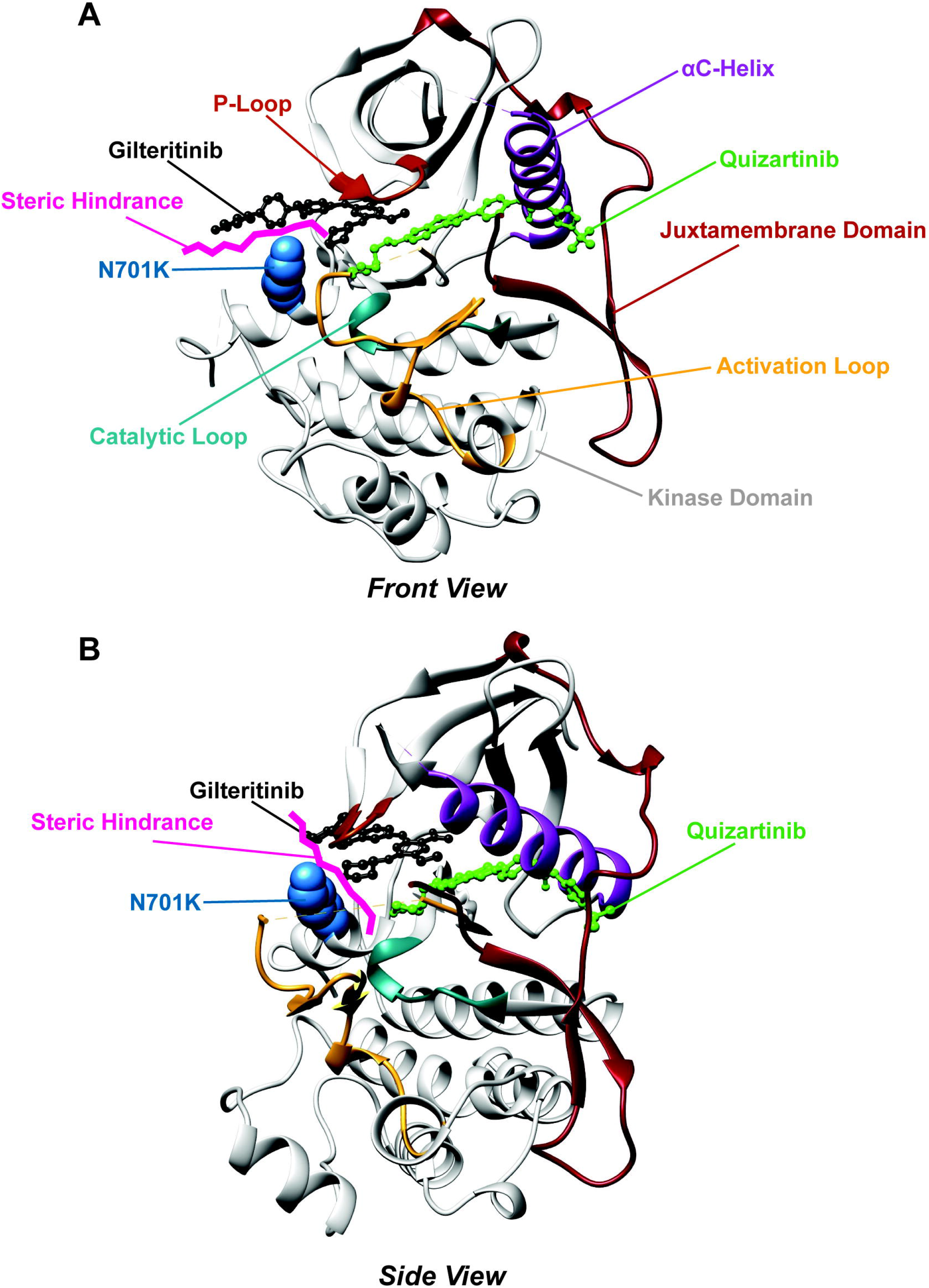

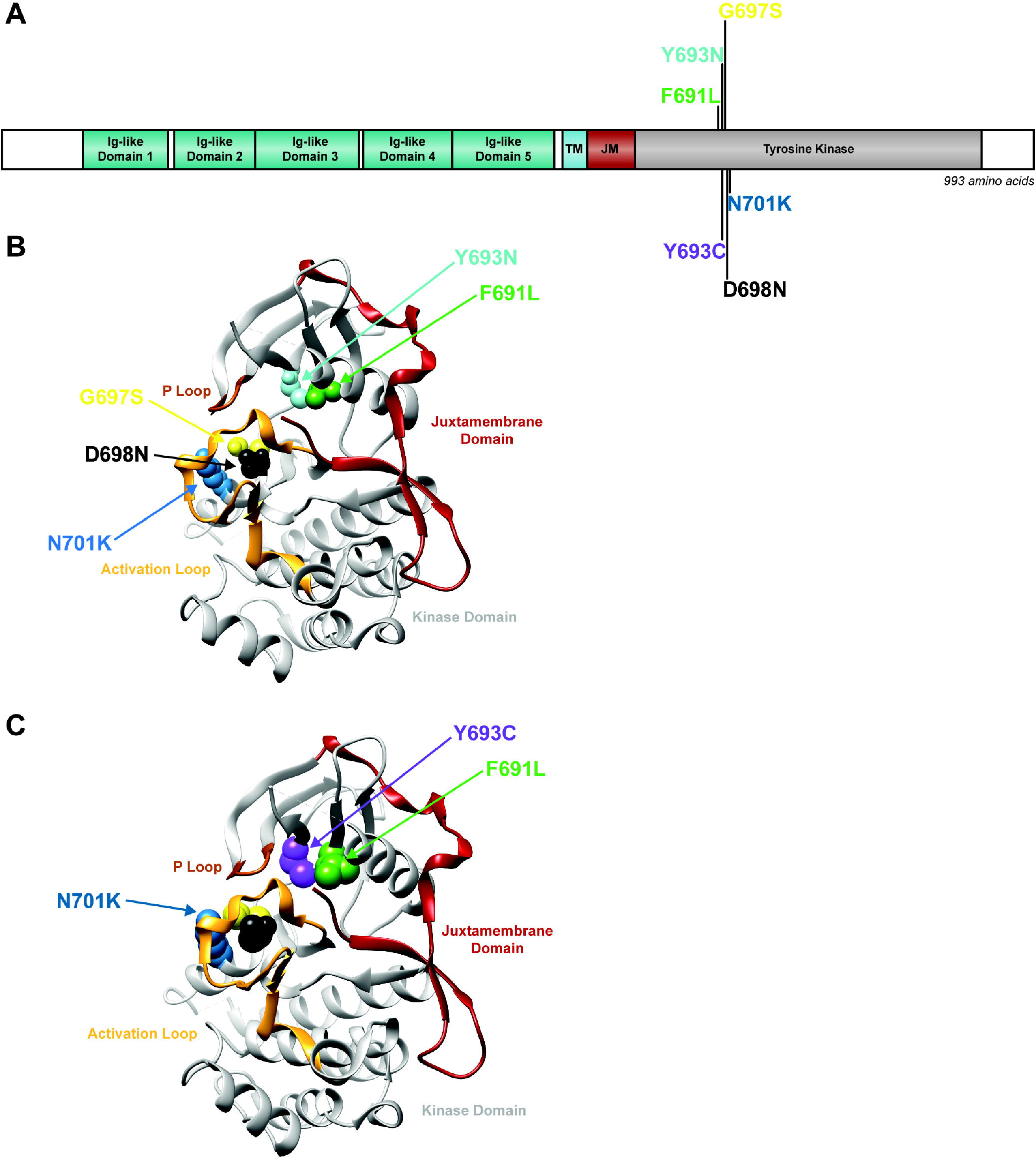

