## Supplemental Methods & Legends for "A noncanonical FLT3 gatekeeper mutation disrupts gilteritinib binding and confers resistance"

### Supplemental Methods, Figure Legends, & Tables

#### TABLE OF CONTENTS:

**SUPPLEMENTAL MATERIALS & METHODS**…………………………………………...…………..2-5

**SUPPLEMENTAL FIGURE LEGENDS**……………………………………………………….……....6

**SUPPLEMENTAL TABLE LEGENDS**……………………………………………………………...…6

**REFERENCES**……………………………………………………………………………...………….7

#### MATERIALS & METHODS:

##### *Generation of early (ligand-dependent) and late (ligand-independent) gilteritinib resistant cultures*

Human MOLM14 cells were generously provided by Dr. Yoshinobu Matsuo (Fujisaki Cell Center, Hayashibara Biochemical Labs, Okayama, Japan). Human MV4;11 cells were purchased from ATCC (Manassas, VA). Both cell lines were grown in RPMI (Life Technologies Inc., Carlsbad, CA) supplemented with 10% FBS (Atlanta Biologicals, Flowery Branch, GA), 2% L-glutamine, 1% penicillin/streptomycin (Life Technologies Inc.), and 0.1% amphotericin B (HyClone, South Logan, UT). Authentication was performed on all cell lines used in this study at the OHSU DNA Services Core facility.

To establish resistant cultures, 10 million MOLM14 or MV4;11 cells were treated with 100 nM of gilteritinib (Selleck Chemicals, Houston, TX) in media alone (N = 4) or in media supplemented with 10 ng/mL of FGF2 (N = 4) or FLT3 ligand (N = 4, FL; PeproTech Inc., Rocky Hill, NJ). All cultures were maintained in 10 mL of media. Every 2 or 3 days, recombinant ligands and gilteritinib were replaced and cell viability was evaluated using the Guava personal flow cytometer (Millipore Inc., Burlington, MA). Following ligand withdrawal, gilteritinib and media were similarly replenished and viability was monitored every 2 to 3 days. All cell lines were tested for mycoplasma on a monthly schedule.

##### *Whole Exome Sequencing*

Genomic DNA for all cell lines was extracted using the DNeasy Blood & Tissue Kit (Qiagen Inc., Germantown, MD) according to the manufacturer’s protocol. High throughput sequencing was performed on all MOLM14 and MV4;11 parental and late/ligand-independent gilteritinib resistant cell lines. Paired-end 100 base reads were generated using the Illumina HiSeq 2500 (Illumina Inc., San Diego, CA) for parental and gilteritinib resistant cell lines after capture with the

Nextera DNA Exome kit v1.2. Each sample was pre-processed using GATK 4.1[1] including alignment to build 37 (GRCh37) of the human genome using BWA[2]. Raw mutations were called for all replicates relative to the parental cell lines using MuTect2. The mutations were annotated using the Variant Effect Predictor v99.1[3]. The final set of mutations were: selected after limiting to those that passed filter, predicted to have a non-synonymous change or indel, seen in less than 1% of GNOMAD[4], and had at least 5 reads with a tumor variant allele frequency (VAF) of greater than 8%. All NRAS and FLT3 mutations were confirmed by Sanger sequencing as previously described (manuscript in review). Sequencing was performed using Eurofins (Louisville, KY) and analyzed using Sequencher and DNASTAR software.

***Site-Directed Mutagenesis***

All FLT3 (F691L, N701K, D835Y) mutations were introduced by site-directed mutagenesis to $FLT3^{WT}$ vector (Addgene, #23895) using the QuikChange II XL kit (Agilent Technologies Inc., Santa Clara, CA) and primers listed in **Supplemental Table 1a**. Mutagenesis primers were designed using the QuikChange Primer Design Program available through Agilent and purchased from Eurofins. Mutated cDNAs were then transferred into a Gateway-modified pBABE-IRES-GFP puro retroviral vector (Cell BioLabs Inc., San Diego, CA) using a Gateway LR Clonase kit (Life Technologies Inc.). Constructs were verified by Sanger sequencing with primers listed in **Supplemental Table 1b**. $FLT3^{D835Y}$ was verified using the M13R primer (5'-caggaaacagctatgacc-3') available via Eurofins.

***Retroviral production, Ba/F3 transduction, and IL-3 withdrawal assay***

HEK 293T/17 cells (ATCC) were cultured in DMEM (Life Technologies Inc.) supplemented with 10% FBS (Atlanta Biologicals), 2% L-glutamine, 1% penicillin/streptomycin (Life Technologies Inc.), and 0.1% amphotericin B (HyClone). To produce murine retrovirus, HEK 293T/17 cells were co-transfected with FuGENE 6 (Promega, Madison, WI), EcoPac helper packaging plasmid, and pBABE FLT3 constructs (wild-type (WT) or mutant). pBABE-IRES-GFP puro was used as an empty vector control. Retroviral supernatants were harvested 48 hours after transfection.

Ba/F3 cells were grown in RPMI (Life Technologies Inc.) supplemented with 10% FBS, 2% L-glutamine, 1% penicillin/streptomycin, 0.1% amphotericin B as stated above, and 15% WEHI conditioned medium (a source of IL-3). Stable Ba/F3 $FLT3^{WT}$ and mutant cell lines were generated by infection of 3 x $10^6$ cells with 1 mL of retroviral supernatant followed by spinoculation with polybrene at 2500 rpm for 90 minutes. Ba/F3 cells infected with pBABE empty, $FLT3^{WT}$, and mutant vectors were then selected using 2 µg/mL puromycin (ThermoFisher Scientific) selection.

Parental Ba/F3 cells and cells expressing empty vector, $FLT3^{WT}$, and mutants were washed three times in RPMI medium with 10% FBS (to remove all traces of IL-3-containing WEHI conditioned medium). Cells were then suspended at a density of 5 x $10^5$ cells per mL and counted on a Guava personal flow cytometer every other day and divided as necessary.

***Small molecule inhibitor studies***

Ba/F3 cell lines were seeded in 384-well plates, incubated with the indicated concentrations of inhibitor for 72 hours, and assessed for cell survival using a methanethiosulfonate (MTS)-based assay (Promega) as previously described[5, 6]. The final concentration of DMSO was ≤0.1% in all wells. Six technical replicates were plated for all Ba/F3 cell lines. Small-molecule inhibitors were purchased from LC Laboratories Inc. and Selleck Chemicals. GraphPad Prism 8 was used to model dose-specific, normalized cell viability values with 4-parameter logistic regression curves to determine $IC_{50}$s.

***Immunoblotting***

Following overnight serum starvation in 0.1% BSA RPMI media, 10-15 million parental, empty vector, and FLT3 mutant Ba/F3 cells were spun down and lysed with 150 μL of Cell Lysis Buffer (Cell Signaling Technologies Inc., Danvers, MA) containing a Complete Mini Protease Inhibitor Cocktail Tablet, Phosphatase Inhibitor Cocktail 2, and Phenylmethanesulfonyl Fluoride (PMSF) solution (Sigma-Aldrich Inc., St Louis, MO) and clarified by centrifugation at 14,000 *g*, 4 °C for 15 minutes. Protein was quantified using a bicinchoninic acid (BCA) assay (ThermoFisher Scientific Inc., Waltham, MA). 50 μg of each protein lysate was loaded on NuPAGE 4-12% Bis-Tris gradient gels (ThermoFisher Scientific Inc.), transferred on Immobilon-FL PVDF membranes (Millipore Inc.), and blocked for 1 hour. Following overnight incubation with primary antibody (**Supplemental Table 2**) at 4 °C, the membranes were washed and probed with fluorescent IRDye 800CW goat anti-rabbit IgG and IRDye 680RD Goat anti-mouse IgG antibodies (1:15,000; LI-COR Biosciences, Lincoln, NE). The membranes were then imaged with the Odyssey Infrared Imaging System (LI-COR Biosciences).

#### SUPPLEMENTAL FIGURE LEGENDS:

**Supplemental Figure 1: $FLT3^{N701K}$ is resistant to type I inhibitors, midostaurin and crenolanib but relatively sensitive to type II inhibitor, quizartinib. A-C.** Six replicates of $FLT3^{WT}$, $FLT3^{ITD}$, $FLT3^{ITD+F701K}$, and $FLT3^{ITD+F691L}$ Ba/F3 cells were plated with a dose gradient (0 – 1000 nM) of type I and II inhibitors, midostaurin (A, type I), crenolanib (B, type I), and quizartinib (C, type II) for 72 hrs. $FLT3^{WT}$ cells were plated in media supplemented with IL-3. Cell viability was determined using a tetrazolamine-based viability assay. Viability is represented as a percentage of the untreated control. The average mean ± SEM is shown.

**Supplemental Figure 2: $FLT3^{N701K}$ sterically hinders binding of type I inhibitor gilteritinib (black).** Binding of type II inhibitor quizartinib (green) is not affected by $FLT3^{N701K}$. This modeling correlates with the drug sensitivity results presented in Supplemental Figure 1. **A.** Front view. **B.** Side view. Ribbon diagrams were adapted from PDB 4RT7[7] (quizartinib) and PDB 6JQR[8] (FLT3 and gilteritinib) and visualized with the UCSF Chimera software[9].

**Supplemental Figure 3: $FLT3^{N701K}$ is found in a region where multiple noncanonical mutations that confer gilteritinib resistance reside. A.** Gene schematic depicts location of all noncanonical gilteritinib-resistant mutations[10]. **B-C.** Noncanonical mutations mapped onto the crystal structure of the FLT3 kinase domain. $FLT3^{N701K}$ is in close proximity to these mutations. Diagram was adapted from PDB 1RJB[11] and visualized with the UCSF Chimera software[9].

#### SUPPLEMENTAL TABLE LEGENDS:

**Supplemental Table 1:** List of all site-directed mutagenesis (a) and Sanger sequencing (b) primers used in this study.

**Supplemental Table 2:** List of all antibodies used in this study.
